# Bayesian *p*-curve Mixture Models as a Tool to Dissociate Effect Size and Effect Prevalence

**DOI:** 10.1101/2024.07.31.606048

**Authors:** John P. Veillette, Howard C. Nusbaum

## Abstract

Much research in the behavioral sciences aims to characterize the “typical” person. A statistically significant group-averaged effect size is often interpreted as evidence that the typical person shows an effect, but that is only true under certain distributional assumptions for which explicit evidence is rarely presented. Mean effect size varies with both *within-participant effect size* and *population prevalence* (proportion of population showing effect). Few studies consider how prevalence affects mean effect size estimates and existing estimators of prevalence are, conversely, confounded by uncertainty about effect size. We introduce a widely applicable Bayesian method, the *p*-curve mixture model, that jointly estimates prevalence and effect size by probabilistically clustering participant-level data based on their likelihood under a null distribution. Our approach, for which we provide a software tool, outperforms existing prevalence estimation methods when effect size is uncertain and is sensitive to differences in prevalence or effect size across groups or conditions.

## Introduction

Many psychology and cognitive neuroscience studies support their claims by showing that there is a statistically significant difference in the mean size of some effect – say, a difference in some variable as the result of an intervention – between two or more conditions or groups. A statistically significant group-level effect is often interpreted to signify an effect is typical in the population, and a difference in the average effect size is often interpreted as a change in the effect size’s magnitude between groups or conditions. This interpretation follows from a set of typical, though usually unstated, assumptions: (1) a group condition mean characterizes the condition response of a “group-representative participant,” (2) measurements from actual participants represent noise around this central tendency, and (3) participants who are very far from the mean are “outliers” who may safely (or even should) be discarded from analyses because they have systematic characteristic differences from the representative hypothetical participant being investigated. In reality, however, a group mean may significantly differ from its null value even when a minority of individual participants show an effect ^1^. Concretely, if some participants come from a null distribution with expectation *E*[*X*|*H*_0_] and others an alternative distribution (where participants show some effect) with *E*[*X*|*H*_1_] ≠ *E*[*X*|*H*_0_], then the expected mean of the population distribution will differ from the null *E*[*X*|*H*_0_] whenever the prevalence of the effect is *nonzero,* not just when it is typical (see Fig. 1). Claims concerning the typicality of an effect should be supported by direct estimation of the effect’s population prevalence, not by a test of group means.

**Figure 1:**
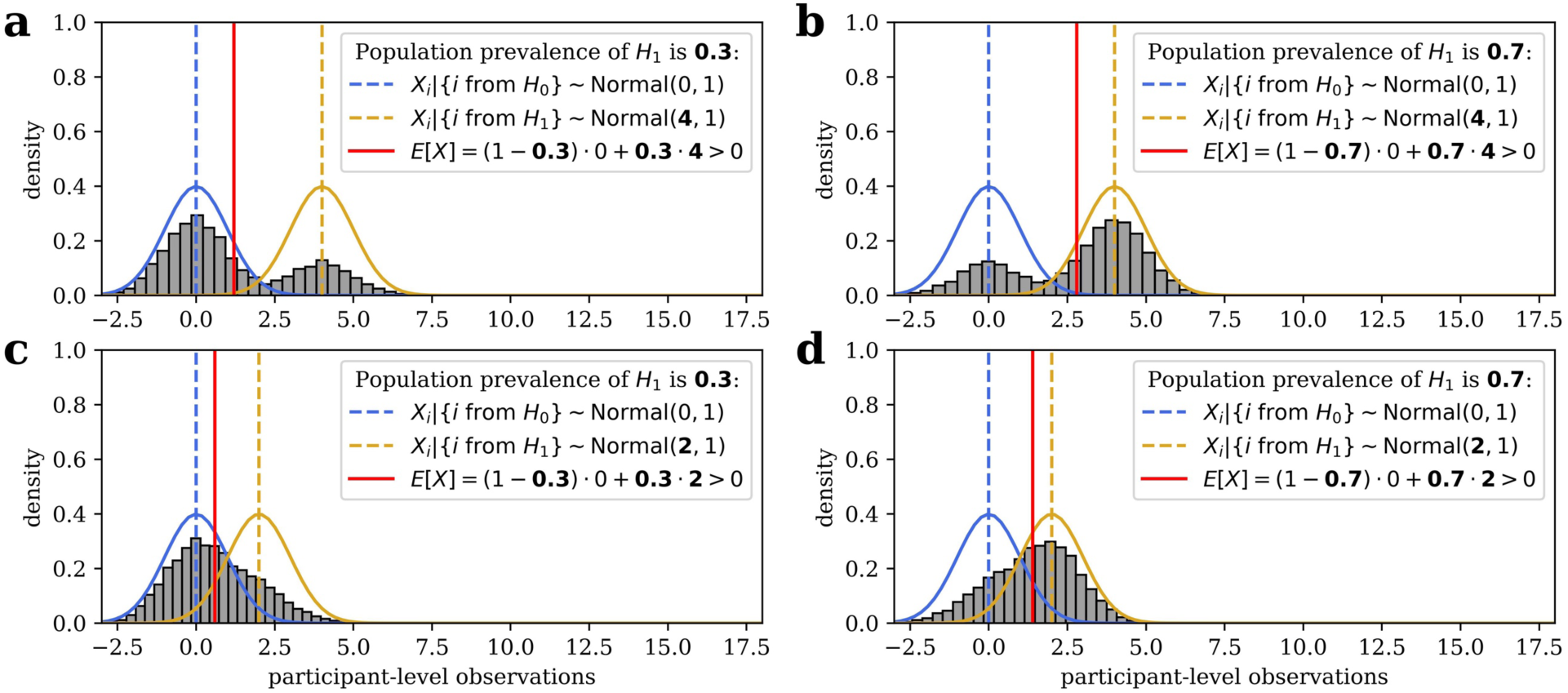
A statistically significant group mean does not imply all, or even most, participants in population show an effect. (a) For example, this histogram shows 10,000 simulated draws from population in which 30% of participants show an effect with a group-level effect size of Cohen’s *d* = 4, and all other participants show no effect or Cohen’s *d* = 0. The two underlying sampling distributions are overlaid with their means, as well as the mean/expected value of the full population. A null hypothesis significance test with this data would rightly reject the null hypothesis that the population mean is equal to zero. (b) For illustration, a similar simulation is shown where prevalence of effect is 70% of population. (c-d) Same as the top panels, but the effect size among participants who show the effect has been reduced to Cohen’s *d* = 2. While still a very large effect size by conventional standards, the histogram is no longer clearly bimodal. Still, population means differ from zero even when only a minority of participants show the effect. Because such a result is not a false positive – the true population mean does, in fact, differ from zero – collecting more data makes it more, not less, likely that a group-level null hypothesis test would reject the null hypothesis as a result of a low prevalence effect.

*Population prevalence* refers to the proportion of a population (or sample from that population) demonstrating an effect (see Table 1). In other words, the mean effect size in a population is, notably, only equal to the *within-participant* effect size if the *population prevalence* of the effect is 100%. If an effect is heterogenous – some people show it and some do not – then the sample mean will be “watered down” relative to the within-participant effect size, and will thus be representative of neither the subgroup that shows the effect nor that which does not. In an experiment, the mean effect size could vary across groups or conditions because the within-participant effect size differs, but it could also vary if the *proportion of people who show the effect* differs.

**Table 1:** Glossary.

| Terms | Definitions |
| --- | --- |
| Group mean effect size<br>Average effect size | A quantitative measure of the strength of an effect obtained by aggregating across all participants within a sample. Depending on the setting, an “effect” may refer to a difference between experimental conditions, association between variables, or measure of task performance. A group mean effect size is a function of both the proportion (prevalence) of participants who show the effect and the within-participant effect size among those participants. |
| Within-participant effect size | A measure of the strength of an effect shown by a single participant, such as estimated from their trial-level data. Averaging the within-participant effect sizes across all participants would yield a group mean effect size. |
| Within-participant power | The sensitivity of a null hypothesis significance test to detect a true effect, based on the trial-level data from a single subject. This is a function of the within-participant effect size, within-participant sample size (i.e. trial count), and an arbitrarily chosen significance level.<br>Effect sizes estimated using *p*-curve mixtures are unitless, so it may be more interpretable to discuss within-participant power instead of within-participant effect size, even though the use of *p*-curve mixture models does not actually entail conducting a null hypothesis test at the participant-level. |
| Population prevalence | The proportion of participants in the population who show some effect, as opposed to no effect or an effect size essentially equal to some null value (e.g. zero difference between conditions or chance accuracy). |
| Bayesian mixture model | A general method of modeling data in which observations are assumed to belong to multiple distinct groups. Observations (i.e. participants) are probabilistically clustered, such that the proportion of observations belonging to each group and the distribution of observations within groups can be inferred simultaneously. |
| *p*-curve | A probability density of independent *p*-values obtained from repeated (for our purposes, identical within-participant) experiments. If *p*-values are calculated properly and assumptions are met, then *p*-curves should be equivalent to the standard uniform distribution for observations drawn from the null distribution and increasingly skewed for larger within-participant power. |
| *p*-curve mixture model | A reparameterization of a two-group Bayesian mixture model such that participant-level observations can be provided as *p*-values rather than in their original units. Observations are modeled as a mixture between a uniform distribution and a skewed *p*-curve. The proportion of participants the model assigns to the *p*-curve is an estimate of the population prevalence. The area under the *p*-curve (with skewness inferred from data) up to some chosen threshold is an estimate of the within-participant power of the experiment from which data were generated. |

This distinction bears important implications. For example, an intervention that has a strong within-participant effect, even if it only applies to a modest subpopulation, could still have useful practical or clinical applications. Conversely, an intervention that achieves an equivalent group mean effect size when averaging over a negligible within-participant effect that is widely present in the population may be less useful in practice – but also more likely to replicate in a new sample, as a researcher does not have to be “lucky” in sampling participants from the selected subgroup. While may be tempting to assume such situations are exceptions, the strong correlation between group mean effect size and the between-study heterogeneity of that effect among preregistered replications indicates the largest effects are actually the most likely to vary (at least in size) across participants ^2^, and even some famously robust patterns of behavior such as in the Simon task have been found not to be universal ^3^. Indeed, there is a growing recognition across fields of behavioral research that effects measured and mechanistic models fit at the group-level often deviate markedly from what is observed in any single participant ^4–8^. As such, Bryan and colleagues have recently argued that the antidote to the reproducibility crisis in the behavioral sciences is a “heterogeneity revolution,” in which systematic approaches to sampling and to quantifying population heterogeneity are adopted ^9^. While we agree with Bryan et al. (2021) that researchers should more often adopt sampling plans that explicitly aim to capture population heterogeneity when measuring already-established effects, much research aims merely to establish the existence of novel effects predicted by theories. Even for the latter sort of research, it would be highly informative for empirical studies to estimate and report the proportion of the (sampled) population to which observed effects can be expected to apply, as well as useful quantities such as within-participant effect size or power estimates, both of which can inform sampling approaches for future confirmatory or applied research.

Currently, standard approaches to dealing with population heterogeneity are likely to exacerbate this problem; for instance, outlier removal aims to eliminate subgroup differences before computing descriptive or inferential statistics. This approach yields sample mean effect size estimates that neither are good estimates of the true population mean, as they exclude parts of the population, nor capture real heterogeneity. A better approach would be to jointly estimate the population prevalence of observed effects and the within-participant effect size for those to whom the effect applies, so as to quantify heterogeneity instead of artificially removing it. Moreover, a systematic difference in the prevalence of an effect between two populations (i.e. between-group difference) may be of scientific interest; for example, as psychological differences between people from Western and non-Western cultures have become increasingly well-documented, it would be fruitful to dissociate whether such differences reflect the prevalence or size of effects, possibilities that suggest distinct theoretical explanations ^10,11^. Moreover, one might wish to estimate the difference in the prevalence of two effects in the same population (i.e. within-group difference) or the conditional probability/prevalence of one effect given that a person shows a different effect – which may be used as evidence that two behavioral and/or neural phenotypes are driven by the same latent trait.

When the ground truth status of individuals is known, estimating prevalence is simply a matter of computing the sample proportion of individuals showing the effect. In practice, however, we must deal with the possibility that some participants may show an effect in reality, but we failed to detect it in our experiment. In such cases, sample proportions are systematically lower than the population prevalence, so frequentist approaches to prevalence inference have primarily aimed to put a lower bound on population prevalence, rather than an estimate ^12,13^. Without such an estimate, however, one cannot easily compare groups or conditions. Ince and colleagues proposed a simple but powerful Bayesian method in which, given the binary outcomes of null hypothesis significance tests (NHST) conducted within each participant, models the incidence of a significant within-participant test as a Binomial distribution as a function of the population prevalence and within-participant power ^14^. Not only does this approach allow prevalence to be compared across groups or conditions, but it is applicable whenever a *p*-value can be computed for each participant. However, estimates are sensitive to the arbitrary significance level of the within-participant test, and as we show here, the approach cannot be fruitfully extended to settings in which the within-participant power/effect size is unknown or may vary across groups or conditions. That is, the Binomial model cannot simultaneously estimate both population prevalence of an effect and the within-participant effect size.

In our past work, we have dealt with this issue by using Bayesian mixture modeling ^15,16^. This approach models the distribution of the participant-level observations as a mixture of two or more subgroup distributions, allowing one to infer the parameters of these distributions and the proportion of participants that belong to each subgroup simultaneously. However, using this approach for prevalence estimation requires constructing an appropriate Bayesian model of the data, which may require substantial expertise or might even be intractable. In the present work, we aim to combine the advantages of mixture modeling with the broad applicability of the Binomial model. A “*p*-curve mixture model” simultaneously infers the population prevalence and the (relative) within-participant effect size from the unthresholded *p-*values of within-participant null hypothesis significance tests. Thus, it can be uniformly applied to any study in which a significance test can be performed per-participant. Since it does not require assuming a fixed power *a priori* as do previous methods, our method can be applied to datasets that were not originally collected with within-participant statistics in mind; this wide applicability affords the opportunity for researchers to estimate population heterogeneity using existing data and experimental designs.

## Methods and Data

### Bayesian mixture models for prevalence estimation

We will first describe how Bayesian mixture models can be used to estimate prevalence and effect size generally, before moving on to *p*-curve mixture models which are a special case. An “effect” here may, for example, refer to that of a treatment of intervention, or to a quantifiable behavioral characteristic/skill – such as a performance measure on a task – that can be compared to a meaningful null (“no effect”) distribution, as in the example in this section.

The general Bayesian approach to statistics is to first describe a generative model of one’s data – that is, to specify the likelihood of the data given some parameters – and then use Bayes’ rule to infer plausible values for the parameters given the observed data. In a Bayesian mixture model, this generative model entails that an observation can come from any one of at least two distributions; thus, the marginal likelihood of the data is a mixture of the likelihoods of the component distributions, weighted by the probabilities of an observation coming from each component. In most applications of Bayesian Mixture models, the parameters of *all* component distributions (e.g. their mean and scale) and the relative probabilities of each component distribution are all estimated during model fitting. This is what most mixture modeling software packages, Bayesian or frequentist, implement out-of-the-box ^17^. Mixture models, Bayesian or otherwise, have often been used as a sort of probabilistic clustering technique to infer the presence of latent subgroups within a large-sample datasets ^18,19^, or to infer effect heterogeneity across studies in meta-analyses ^7^. However, mixture models, as have other sorts of hierarchal models have been used to assess heterogeneity in the literature ^20^, can be applied to achieve more specific inferential goals by placing constraints on the shape of the component distributions ^21^.

When one component distribution is constrained such that one component distribution is a null *H*_0_ distribution – the likelihood of a participants’ data if there is no effect – and the other distribution is an alternative *H*_1_ distribution which ideally incorporates prior information about the range of plausible effect sizes under the alternative hypothesis, then the probability γ assigned to the *H*_1_ distribution (i.e. the *H*_0_ or *H*_1_ group membership of each participant is modeled as a Bernoulli(γ) distributed random variable) is an estimate of *H*_1_’s prevalence in the sampled population. γ can be given a Uniform(0,1) or Beta prior distribution.

Once the model and prior distributions are specified, the posterior distribution for each parameter – representing beliefs about plausible parameter values after conditioning on the observed data – is given by Bayes’ rule. A normalizing constant for the posterior distribution (i.e. to make it sum to 1) for a given parameter cannot usually be derived in closed form, but the posterior can be sampled from without knowing the normalizing constant using Markov-Chain Monte Carlo ^22^.

To estimate prevalence with a traditional Bayesian mixture model parametrization, one needs to (a) be able to specify the likelihood of the data under the null hypothesis and (b) under the alternative hypothesis. Moreover, they must (c) know how to program the custom model and approximate posterior distributions for its parameters in a probabilistic programming framework such as *PyMC* or *Stan* that will perform posterior sampling for us ^23,24^. We simply do not always have these ingredients. Often, even the shape of the null distribution is not known a priori – e.g. the same situations where one might use a non-parametric test in the NHST framework. Moreover, it is not practical to assume all researchers have the combination of domain and technical knowledge to implement such an ad hoc statistical model for their own, idiosyncratic dataset. If there were, however, a way to transform observed data from arbitrary distributions into a distribution we always know enough about to model this way – much like users of NHST can almost always resort to a non-parametric test – then researchers would not need to write their own software for each use case. This limitation of traditional Bayesian mixture modeling parametrizations motivates the use of the *p*-curve mixture models, which is a special case of a Bayesian mixture model in which the likelihood of the data is written in terms of the data’s *p*-value rather than in terms of its original units.

### The *p*-curve likelihood

Since *p*-values most commonly appear in the setting of null hypothesis significance testing (NHST), it may be unconventional to think of them as random variables. Indeed, in studies in which only one *p-*value is computed or multiple *p*-values are computed on the same data, thus are not independent observations, it would not be useful to think of them this way. However, when a test is repeated multiple times on *independent* samples, then the independent *p*-values are indeed random variables with a probability distribution. This sort of distribution, called a *p*-curve, has been leveraged to study and correct for the effects of publication bias in the meta-analysis literature^25–27^.

Under the null hypothesis, a *p*-curve is always Uniform(0, 1); this fact is built into the logic of NHST, since the *p*-value can only fall under the significance level α just α proportion of the time for *any* arbitrary α. Under the alternative hypothesis, the *p*-curve becomes increasingly left skewed the larger the effect size or sample size (see Figure 2). One can derive an exact *p*-curve for a given statistical test, such as the *t-*test, as a function of the effect size and sample size ^25^. Usefully, such an approach can be used to recover an effect size merely from repeated *p*-values as has been done in meta-analysis ^26^.

**Figure 2:**
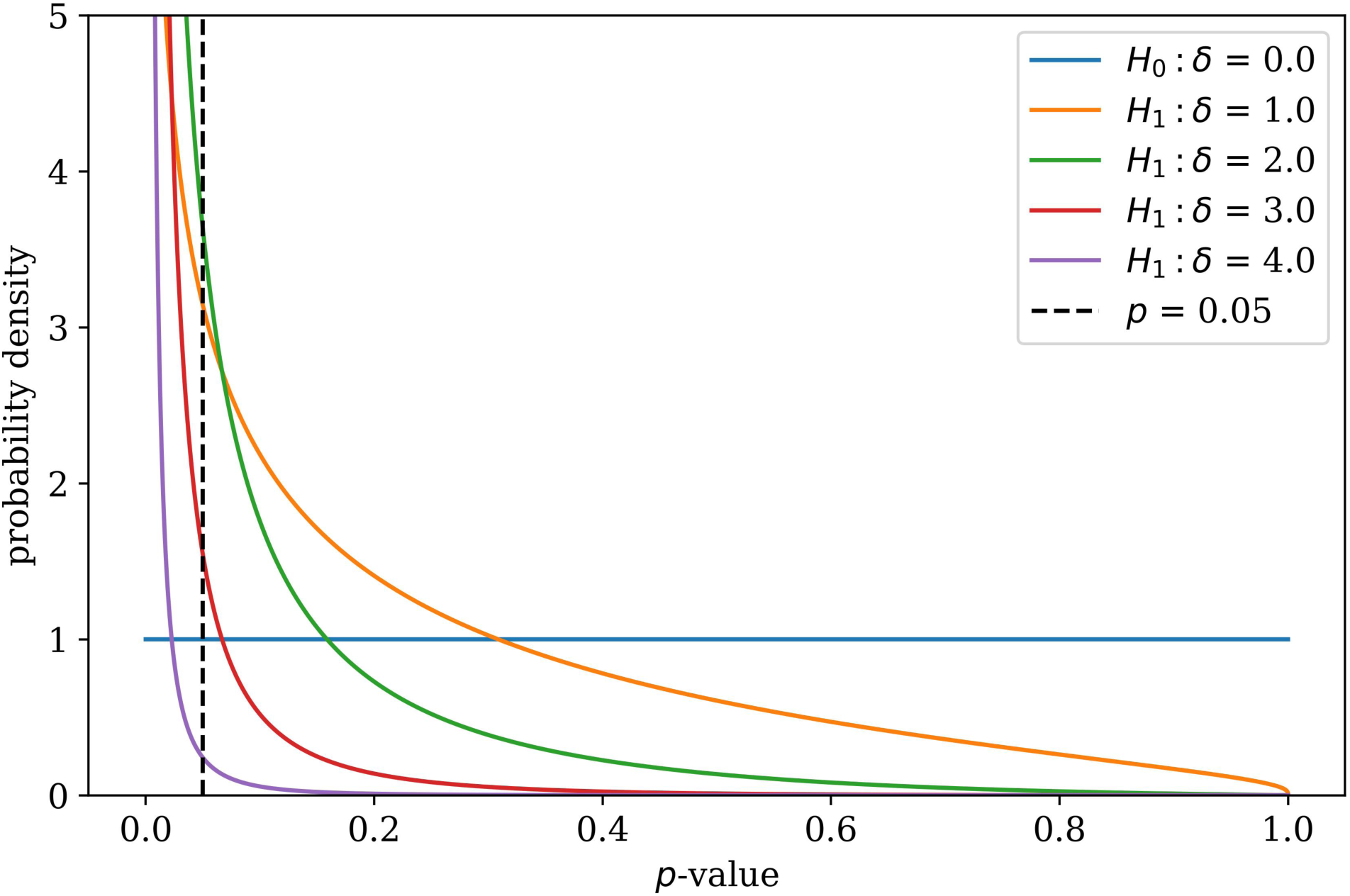
A probability distribution for *p*-values. Describing the likelihood of *p-*values from repeated, independent tests of an effect with size δ, these *p*-curves – specified by Equation 1 – are valid probability distributions that integrate to one over the interval (0, 1). When the null hypothesis is true (δ = 0), the *p*-curve is equivalent to the standard uniform distribution.

However, since the “sample size” is not always well-defined for a within-participant study (e.g. consecutive trials in a task or neuroimaging measurement may not be independent observations), we specify a probability density function *f* for a *p*-curve in Equation 1 that depends on a single effect size parameter δ using the standard normal probability density function φ and cumulative distribution function Φ.

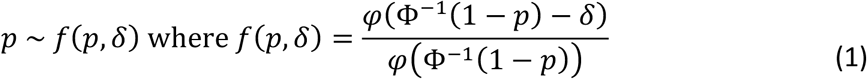

The use of the normal distribution in our specification does not entail that our model will work only with normally distributed data – unless you happen to be using a Z-test with sample size one, in which case this *p*-curve is indeed exact – as we will verify in our simulations. It should be duly noted though that, without a sample size parameter, the “effect size” parameter in our model is abstract/unitless and should only be compared across identical experiments (same number of trials, design, etc.) with an identical method of computing *p*-values (e.g. one- or two-tailed). However, this unitless effect size parameter can be converted back into meaningful units by taking the calculating the area under the *p*-curve distribution up to (i.e. cumulative distribution function at) some *p* = α, which is interpreted as the power of a within-participant NHST at significance level α.

### The *p*-curve mixture model

Now that we have a probability distribution that can describe how *p*-values are distributed given an effect size, we can incorporate it into a mixture model – which can then be applied whenever per-participant *p*-values can be computed using NHST within each participant, regardless of the original distribution of the data. A very similar approach – using *p*-curves as a conditional likelihood in a Bayesian mixture model – has been applied with group-level *p*-values as observed data for the purpose of metanalysis, aiming to estimate the proportion of false positives contaminating published literatures ^28^. As in a more typical mixture model for prevalence estimation, participants show an effect (i.e. *H*_1_ is true) with probability/prevalence γ. If *H*_1_ is true for participant *i*, then their *p*-value is distributed according to *p_i_*∼*f*(*p_i_*, δ), as defined in Equation 1, for some unitless effect size δ. If instead *H*_0_ is true, then *p_i_* ∼*f*(*p_i_*, 0) or equivalently *p_i_* ∼Uniform(0,1).

Before performing Bayesian inference given a set of observed *p*-values, one needs to assign prior distributions to γ and δ. In all simulations and examples in this manuscript, we will use an uninformative γ∼Uniform(0,1) prior for the prevalence of *H*_1_ and a weakly informative prior of δ∼Exponential(2/3). The latter was chosen as it results in a prior 90% highest-density interval for within-participant power (at significance level α = 0.05) of roughly [0.05, 0.95], and thus is relatively close to a uniform prior over within-participant power. Note that a flat prior, instead of an exponential prior, would allow unreasonably high values of δ and would not be “uninformative” in this case. Our exponential prior, per our simulations, performs well across a variety of cases.

### Between-group differences in prevalence or effect size

*p*-curve mixtures can be used to estimate the difference in prevalence and in (unitless) within-participant effect size between two independent groups of participants. This is useful, for example, in a between-participant experimental design with random assignment, or when comparing experimental results between two distinct populations (e.g. men and women, liberal and conservatives, Westerner and Easterners, etc.).

If the groups are independent, and thus group differences can be treated as fixed effects, then it is appropriate to estimate the difference by fitting *p*-curve mixture models to each of the two populations separately, drawing samples from the parameters of both model posteriors (see ***Estimation and Software***), and subtracting the samples between the groups’ models to approximate samples from the posterior of the difference. We use this approach in our EEG simulations, as well as in Example B and its accompanying code notebook.

### Within-group differences in prevalence or effect size

Differences in prevalence and effect size can also be estimated *within* a group, if one has conducted two tests per participant with alternative hypotheses *H*_1_ and *H*_2_. One may find it useful to do so if they have measured the same effect in two within-participant experimental conditions (i.e. an experiment designed to test for an interaction effect) or if they want to assess whether showing one effect makes it more or less likely that the same participant shows some other effect (e.g. whether above-chance performance on one task is more or less prevalent than above-chance performance on another task). It may even be used when a researcher wants to know if participants who pass some manipulation check are more or less likely to show an effect of interest – an alternative to outlier rejection that does not require binary thresholding but is instead uncertainty-weighted.

In this case, unlike in the between-group case, models cannot be estimated separately for each test, since participants are observed in both tests and thus observations are not independent. In this joint model, each participant either shows no effect in either test (denoted *k*_00_ = 1), just the *H*_1_ effect (denoted *k*_10_ = 1), just the *H*_2_ effect (*k*_01_ = 1), or both effects (*k*_11_ = 1). *k*_00_, *k*_10_, *k*_01_, and *k*_11_ are Categorical/Multinomial(1) distributed with probabilities θ_00_, θ_10_, θ_01_, θ_11_, respectively. The prevalence of *H*_1_, then, is γ_1_ = θ_10_ + θ_11_ and of *H*_2_ is γ_2_ = θ_01_ + θ_11_. The conditional probability that *H*_2_ is true for a participant given that *H*_1_ is true is *P*(*H*_2_|*H*_1_) = θ_11_/γ_1_ and vice versa. We place a minimally informative Dirichlet(1, 1, 1, 1) prior (a common default) on the θ’s.

This extended model is actually somewhat difficult to implement, as the discrete latent variables *k* must be marginalized out analytically to ensure robust sampling from the posterior during estimation. We have provided, as for the basic *p*-curve mixture model, a user-friendly wrapper around our optimized implementation (see ***Estimation and Software***).

### Estimation and Software

Posterior distributions for all parameters in a *p*-curve mixture model can be approximated by drawing thousands of samples from the posterior (we use 5,000 in our simulations and examples) using a No-U-Turn sampler or other Markov Chain Monte Carlo sampler ^22^. This procedure would be laborious to program by hand, but fortunately a number of “probabilistic programming” frameworks can perform this sampling for you, which makes the *p*-curve mixture model quite simple to implement for those already familiar with Bayesian modeling ^23,24^; our implementation uses *PyMC* for posterior sampling. Posterior samples can be summarized using the posterior mean/expectation (estimated as the mean of the posterior samples and the 95% (or whatever percent) highest density interval (HDI), which is the smallest interval such that there is a 95% posterior probability the estimated parameter falls within that interval.

We expect most researchers who could benefit from *p-*curve mixture modeling may not be familiar with probabilistic programming or Bayesian statistics in general. To this end, we built a user-friendly Python programming interface to our model implementation so that *p-*curve mixture models can be fit and summarized in just a few lines of code, as well as a graphical interface for users who are not familiar with Python programming (see **Code Availability** for details).

### The Binomial model

As a baseline to which to compare *p*-curve mixture models, we also implement a version of Ince et al.’s (2021) Binomial model for estimating population prevalence ^14^, extended to accommodate uncertainty about within-participant power/effect size. In this model, the probability of rejecting the null hypothesis for each participant is π = (1 − γ)α + γ(1 − β), given the prevalence γ, the Type I error rate α, and the Type II error rate β of the within-participant NHST. The observed number of rejections *k* out of *n* participants, then, is *k*∼Binomial(*n*, π).

Ince et al. (2021) note that, if values of α, β are fixed, then π is a deterministic function of γ. Instead of estimating the multilevel model they estimate the posterior distribution of π given observed *k*, which can be calculated analytically if a π∼Beta(1,1) ≅ Uniform(0,1) prior is used for π (due to the “conjugacy” property of the beta and binomial distributions). They then solve for the implied posterior of prevalence γ algebraically. However, the Type II error rate β – a function of the within-participant effect size – is usually not known a priori. Ince et al., then, simply set β = 0 (i.e. within-participant power equals 1), such that their prevalence estimates are, in fact, a lower bound on the true prevalence – which is only a tight lower bound when statistical power is effectively 100%.

The Binomial model can easily be (though, as we show in **Validation** should not be) extended to the setting where β is unknown by putting uniform priors on both β and γ directly, then approximating their joint posterior distribution using Markov Chain Monte Carlo ^22^, as we do for our *p*-curve model. Intuitively, it seems like this would yield the same benefits as our *p*-curve mixture model – namely, simultaneous estimation of population prevalence and within-participant power/effect size. However, this is not the case; the fact that the model considers the binary outcomes of the participant-level NHST as observations instead of the unthresholded *p*-values results in a likelihood in which an observed 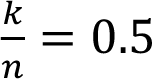 could be (almost) equivalently well explained “ by γ = 1, β = 0.5, by γ = 0.5, β = 0, resulting in joint posterior distributions of γ, β that are near-symmetric about the line γ = 1 − β. As a result, the posterior expectation of γ and of (1 − β) both tend toward the same value – the average of the true power and prevalence – as *n* increases, rather than tending toward their respective ground truths. As we will illustrate in Example C, changing the significance level α can also change the binary outcomes of the within-participant NHSTs thus changing the observed *k*, which can lead to drastically different posterior estimates when *n* is small. Ideally, an estimator would not be so sensitive to the value of an arbitrary parameter.

The extended Binomial model can be used for between-group comparisons of prevalence or within-participant power in the same manner we described for the *p*-curve mixture model (see ***Between-group differences***), and we compared the sensitivity of that approach using the two models in our simulations. Ince et al. (2021) also describe an analog approach to estimating within-group differences, but we did not implement that version of their model here.

### Bayesian model comparison

An advantage of framing the problem of estimating prevalence and effect size as inferring on the parameters of a generative model is that the model can be compared using the full suite of Bayesian model comparison tools. In particular, the null (all *H*_0_) model, the alternative (all *H*_1_), and the mixture (some participants *H*_0_ and others *H*_1_) can all be compared, as all of these models are described by *p*-curve likelihoods. Critically, this allows a researcher to perform Bayesian hypothesis testing to assess whether the population distribution of an effect is indeed heterogenous. Notably, since the prevalence is a proportion, the posterior HDI will never contain 0 nor 1; thus, the mere fact the HDI excludes these values cannot be used as evidence for population heterogeneity. Only model comparison can be used to support claims about the presence of absence of population heterogeneity when the HDI is close to these boundaries.

One approach to Bayesian model comparison is to estimate the leave-one-(participant)-out cross-validated posterior likelihood of the data, which can be computed – with an uncertainty estimate – efficiently using Pareto-smoothed importance sampling or PSIS ^29,30^. Intuitively, the model under which the likelihood of the held-out data is the highest is also the most likely to explain new data. “Weights” can be computed for each model which are, roughly speaking, a probability that each model will predict new observations better than the other models ^29^. Because this method of model comparison is both computationally efficient and relatively numerically stable, we have incorporated it into the *p2prev* package so that users can estimate leave-one-out likelihoods and model weights with just a couple of lines of code.

However, the PSIS approach’s emphasis on prediction may not always be desirable. If, for example, the true prevalence of an effect is zero, a *p*-curve mixture model should estimate a posterior prevalence near zero, resulting in posterior predictions quite similar to those of the *H*_0_-only model. PSIS may not differentiate between these edge cases as well as Bayes Factors, which quantify how much the prior odds on which model is *correct* (rather than most predictive) should be updated given the observed data. We estimate Bayes Factors by nesting the *H*_0_-only, *H*_1_-only, and *H*_0_/*H*_1_-mixture models within a single, hierarchical model with a Categorical prior over models ^31^. This nested model is numerically difficult to sample from due to the discrete latent variable, which can require troubleshooting the sampler; consequently, we provide a code example for estimating Bayes Factors in *PyMC* (corresponding to Example C) but do not build this method into the *p2prev* package itself as we cannot pick default sampler settings that will work in all cases. We also note that there are other methods for computing Bayes Factors for which we do not currently provide examples but may sample more robustly than our nested model approach in some cases. Specifically, marginal likelihoods of models (and thus the ratio of those likelihoods, a Bayes Factor) can be estimated using Sequential Monte Carlo approaches ^32,33^ among which bridge sampling seems particularly promising ^34^.

It should be noted that Bayes factors can be sensitive to the effect size (δ) prior, so researchers should ensure their choice of δ prior does not unduly affect their results if they deviate from validated default priors.

In cases where the population distribution is indeed heterogenous, the *H*_0_/*H*_1_-mixture model should be sufficiently distinguishable from the *H*_0_-only and *H*_1_-only models using the PSIS method. In cases where the sample size is very small or if one wishes to support a claim that either the *H*_0_-only or the *H*_1_-only model is true, we recommend using Bayes Factors instead as predictive criteria may not discriminate the models. In addition, for an approach that is agnostic to the effect size under *H*_1_, researchers can also test the *H*_0_-only model in the frequentist framework by combining *p*-values across participants into a “global” *p*-value using Fisher’s method, Simes method, or related approaches ^35^.

## Validation

### Prevalence estimation benchmark

In this set of simulations, we aimed to benchmark the prevalence estimation performance of the *p*-curve mixture model and of the Binomial model across various sample sizes in a setting in which within-participant statistical power to detect an effect using NHST was less than 100%. To this end, we simulated (1) a setting in which within-participant power was low (∼0.6) but prevalence was high (∼.95) and (2) a setting in which those were flipped and power was high (∼.95%) and prevalence was low (∼0.6). We performed 1,000 simulations at every sample size from *n* = 3 to *n* = 60, spaced by 3.

We also wanted to stress-test the *p*-curve mixture model by introducing some everyday violations of its assumptions. The way we have specified the *p*-curve mixture model is such that all participants for which *H*_1_ are modeled as having the same effect size δ. We intend this to be interpreted as the “average effect size given *H*_1_,” but we would like to verify that our specification can tolerate mild over-dispersion of effect size. Consequently, we simulated 50 trials from participants with within-participant classification accuracies, for participants in which *H*_1_ was true, were drawn from a distribution with (1) mean 0.65 ± 0.01 SD in the low-power setting and (2) mean 0.75 ± 0.01 SD in the high-power setting. We computed *p*-values for each participant using a permutation test to show that the within-participant NHST used to generate *p*-values does not need to make normality assumptions or even be parametric, despite the presence of the normal density function in Equation 1. This choice is also motivated by the fact that it is common to evaluate classification performance using permutation tests when one has estimated out-of-sample accuracies by cross-validation, such as in multivoxel pattern analysis in neuroimaging where it is assumed the assumptions of the binomial test are violated ^36^.

On each simulation, we compute the posterior expectation of the population prevalence and the width of the 95% HDI (smaller means less uncertainty) under both the *p*-curve and Binomial models, and we compute the frequentist false coverage rate of the HDI at each sample size, which is the proportion of simulations in which the HDI contained the true prevalence. While Bayesian HDIs do not provide coverage rate guarantees like frequentist confidence intervals, it is nonetheless a desirable property when a 95% HDI achieves a false coverage rate near or less than 5% in practice, which can be evaluated by simulation ^37^.

Like essentially all Bayesian estimators, the *p*-curve mixture model’s posterior mean is a biased estimator; it is heavily influence by the prior distribution when the sample size is small but gets closer to the true prevalence as *n* increases (see Figure 3a). The Binomial prevalence model ^14^, however, is unable to distinguish between population prevalence and within-participant power, so its posterior mean converges near the average of the two as the number of participants increases.

**Figure 3:**
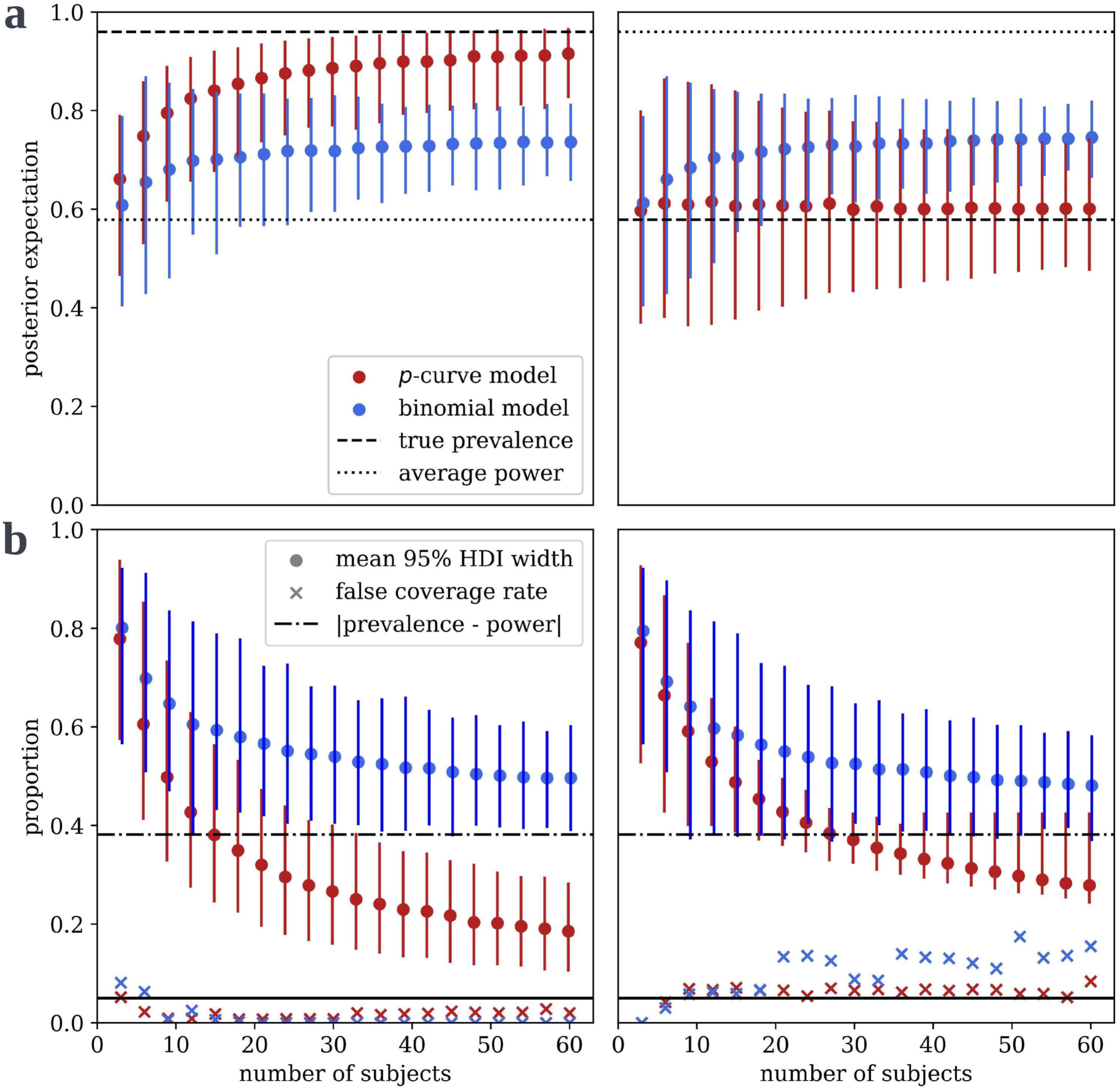
*p*-curve mixture model outperforms state-of-the-art when within-participant power is not known. In all panels, dots reflect the mean across all simulations and bars contain the values from 95% of simulations. (a) The *p*-curve mixture model’s expected population prevalence (a.k.a. posterior mean) gets closer to the true prevalence as it observes more data, but the Binomial model cannot differentiate between prevalence and within-participant power, so its posterior mean converges somewhere between the two. (b) The *p*-curve’s highest density interval (HDI), which contains 95% of posterior probability, shrinks as the model is given more data, reflecting greater certainty, but the Binomial model’s HDI width is lower-bounded by the difference between power and prevalence. The *p*-curve mixture model maintains a false coverage rate (proportion of simulations in which the true prevalence is not contained in the HDI) comparable to that of a frequentist confidence interval.

The *p*-curve mixture model gets (justifiably) less uncertain when it has more data; that is, the interval that contains 95% of the posterior probability (95% HDI) shrinks as the number of participants gets larger (see Figure 3b). Conversely, as the Binomial model cannot distinguish between high-prevalence/low-power and low-prevalence/high-power, its HDI is almost never smaller than the difference between power and prevalence. Thus, after some time, additional data ceases to be useful to the Binomial model, but the *p*-curve mixture model continues to learn from new data. Notably, the *p*-curve mixture model, though it does not come with a false-coverage rate guarantee like a frequentist method would, yields 95% HDIs that achieve empirically strong false coverage rates near or below 5% much like a frequentist 95% confidence interval.

### Robustness to Heterogeneity

Using the same simulation procedure described above with fixed prevalence of 0.4 and mean participant-level accuracy of 0.65, we performed 1,000 simulations at each of 20 ascending levels of between-participant effect size heterogeneity ranging from 0.65 ± 0.005 SD to 0.65 ± 0.1 SD. Again, we estimated the false coverage rate at each heterogeneity level as the proportion of those 1,000 simulations in which the 95% posterior HDI did not contain the true prevalence.

Across all levels of between-participant heterogeneity tested – distributions of participant-level effect sizes visualized in Figure 4a – the false coverage rate of the 95% posterior HDI remained below 5% (see Fig. 4b). One might speculate that this apparent robustness arises from the functional form of the *p*-curve. Looking at Figure 2, it is apparent that the average of the *p*-curves for two effect sizes δ_1_ and δ_2_ is likely to be very close to the *p*-curve for some intermediate effect size in the range (δ_1_, δ_2_). Since our model only uses an approximate *p*-curve to begin with, the fidelity of the approximation may not be particularly different between the *p*-curve for an effect shared by *n* non-heterogenous observations and a *p*-curve obtained by marginalizing over the *p*-curves corresponding to the varying effect sizes of *n* heterogenous observations.

**Figure 4:**
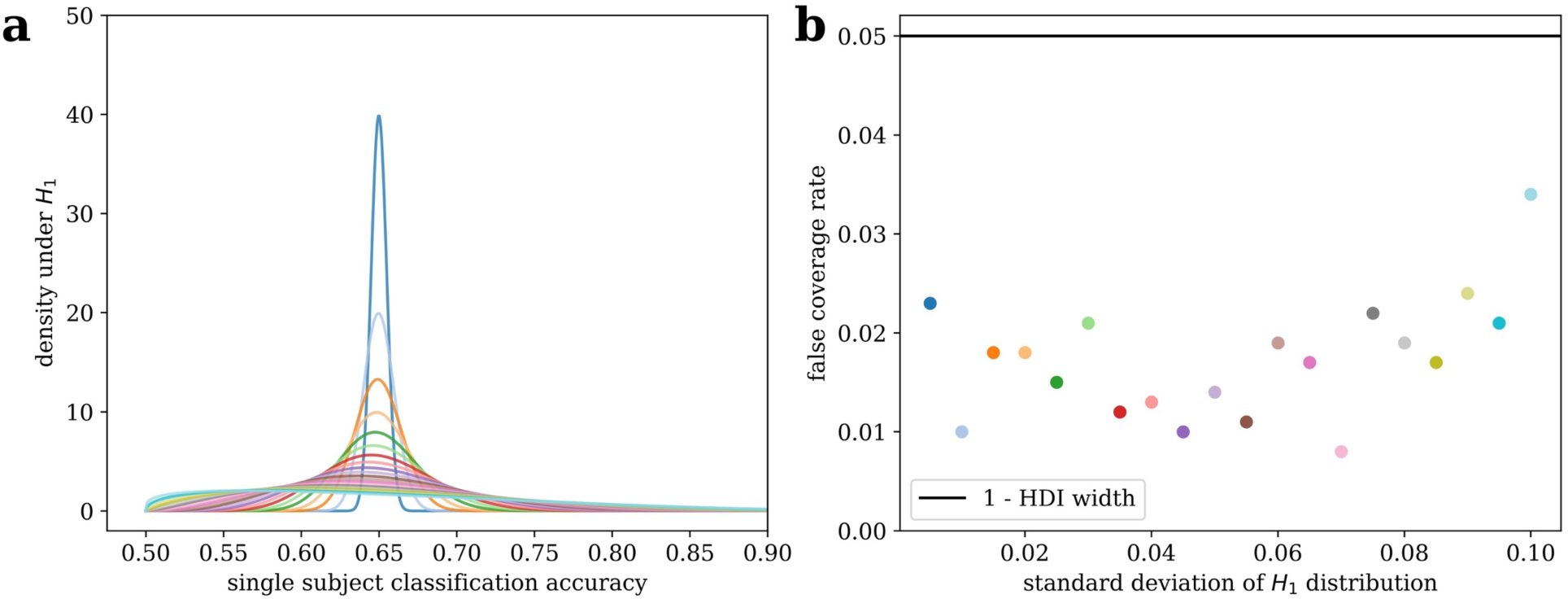
Robustness to Heterogeneity: (a) Distributions of participant-level accuracies under the alternative hypothesis, at various levels of between-participant heterogeneity. The mean of each distribution is 0.65. (b) False coverage rates of a 95% posterior highest-density interval (HDI) over 1,000 simulations with a ground-truth population prevalence of 0.4.

### Sensitivity to between- and within-group differences in EEG data

In this set of simulations, we aimed to assess the sensitivity of the *p*-curve model for detecting between- and within-group differences in population prevalence and in within-participant effect size with realistically noisy data. We compare the *p*-curve model’s sensitivity to that of group-level null hypothesis significance testing, though NHST does not differentiate between prevalence and within-participant effect size contributions to detected changes in the group mean effect size. In the between-group simulation, we also compared the model’s ability to differentiate between changes in prevalence and effect size to that of the Binomial model.

We simulated electroencephalographic (EEG) event related potential (ERP) data by adapting the simulation code used by Sassenhagen and Draschkow, in which a simulated ERP is inserted into background noise that has been cut out of a real EEG recording ^38^. This approach ensures that the noise properties reflect that of realistic EEG data. On each simulation, we simulated 100 trials of EEG data (50 per condition) for each of 30 participants per group. We assigned each participant to *H*_0_ or *H*_1_ with probability equal to the population prevalence γ = 0.45, and we inserted an ERP into one condition of *H*_1_ participants’ data, which we adjusted the size of until the power of a within-participant NHST (at significance level 0.05) was also approximately 0.45. As is the gold-standard in the EEG literature, we computed within-participant *p*-values with an independent-samples cluster-based permutation test which tests for an effect anywhere across all electrodes and timepoints while controlling the familywise error rate ^39^. Naturally, the cluster-level test statistic in this NHST does not have a simple distribution under the null hypothesis – EEG data is participant to substantial noise that is autocorrelated across both electrodes and time – so this is meant to be a strong demonstration of the power of *p*-curve mixture models where it would be impractical to construct a parametric Bayesian mixture model.

In each between-participant simulation, we simulated another group of 30 participants with either higher *H*_1_ prevalence (0.45➔0.95) or higher within-participant power (0.47➔0.96), which was achieved by increasing the magnitude of the simulated ERP. For each within-participant simulation, we generated another 100 trials per participant with similarly increased power or prevalence for some test *H*_2_, ensuring that *H*_2_ is true for all participants in which *H*_1_ was true. This latter scenario reflects how most ERP studies would be designed; a 2×2 design in which one factor is designed to isolate an ERP component of interest (i.e. compute a difference wave), and another factor (i.e. an experimental manipulation) is meant to induce a change in that ERP component.

We perform 1,000 simulations of each of those four contrasts. We then quantify how sensitive the models’ posterior probability of a prevalence increase across groups/conditions is at correctly identifying prevalence increases without false alarming on power increases and vice versa, using the area under the receiver-operator characteristic curve (AUROC) and the true positive rate at a 5% false positive rate. We also apply a group-level NHST on each simulation, first computing the participants’ difference waves (average of 50 trials in one condition minus the average of the 50 trials in the other condition) and inputting them into an independent samples cluster-based permutation test for the between-group simulations and into a paired cluster-based permutation test for the within-group simulations ^39^.

We found the *p*-curve mixture model was highly sensitive to changes in population prevalence and in within-participant effect size in our simulation (see Figure 5). The model is sensitive to between- or within-group differences in prevalence or in within-participant power without mistaking one for the other (see Figure 5b). Moreover, while significant differences in the group mean effect size are often interpreted as differences in within-participant effect size, NHST was highly sensitive to changes in the population prevalence (see Figure 5c). Actually, *p*-curve models outperformed NHST at detecting changes in within-participant effect size, likely because the *p*-curve model’s effect size estimate isolates those participants who actually show the effect – implicitly serving the function of outlier removal but without imposing an arbitrary threshold.

**Figure 5:**
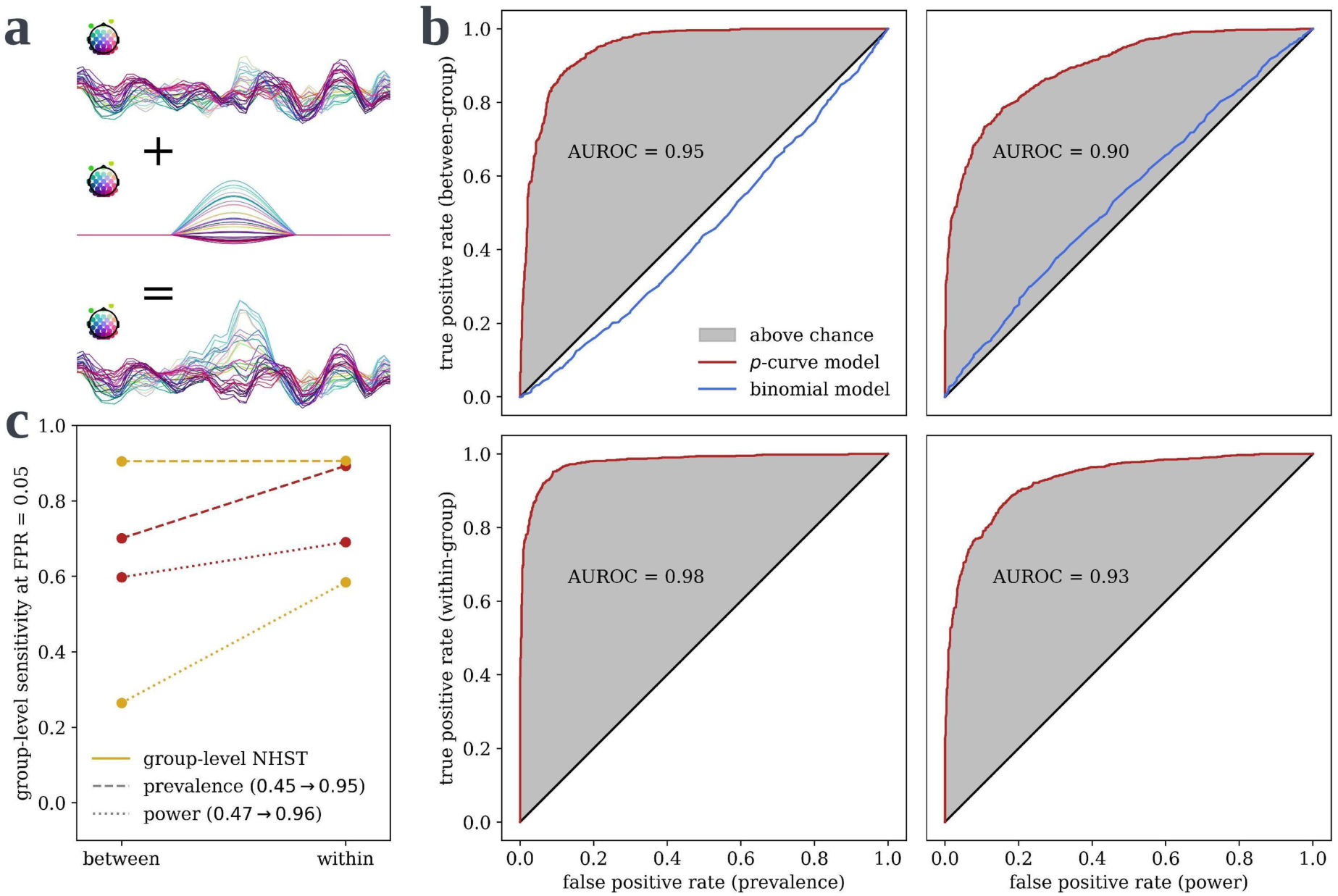
*p*-curve mixture models can detect and discriminate between differences in effect prevalence and effect size between groups or conditions. (a) We simulated EEG data by inserting a simulated evoked response into background noise from a real EEG recording. (b) On each simulation, we simulated two groups of *p*-values for within-participant tests of the evoked response, manipulating either the prevalence of the evoked response, on left (1,000 simulations), or its magnitude in those who show the effect, on right (1,000 simulations). The model was highly sensitive at detecting prevalence or effect size differences between independent groups of participants, on top, or between two within-participant conditions, on bottom. False positives, here, refers to mistaking a prevalence increase for a power/effect size increase or vice versa. (c) The detection rate at the 5% false positive rate is compared to the sensitivity of a group-level NHST with significance level 0.05. In contrast to the normative interpretation of a significant difference in group mean, NHST was highly sensitive to changes in prevalence but less sensitive to changes in effect size than *p*-curve mixtures. In the within-group case, there is no apparent sensitivity cost to using *p*-curve mixtures, which can dissociate between differences in prevalence and within-participant power/effect size.

It is worth noting that, while the *p*-curve mixture models obtained strong sensitivity at a 5% false positive rate (see Figure 5c), the cutoff at which this 5% rate was obtained – that is, the posterior probability of a prevalence or power increase at or above which one would say that there is, in fact, an increase in prevalence or power – is not necessarily 95%. In our simulations, the posterior probability threshold that yielded a 5% false positive rate for differences in effect size was roughly 95% (0.958 and 0.962 for between- and within-groups respectively), but was much lower for differences in prevalence (0.846 and 0.752 respectively). This is not a bad thing; actually, it is a hallmark of high specificity. Our posterior probability of a prevalence difference is insensitive to changes in within-participant effect size, which is what we want. The true Bayesian approach would be to simply report to posterior probabilities, but pragmatically some researchers may indeed have a specific reason to care about frequentist false positive rates. Such a researcher could always take a brute-force approach – at the cost of computation – and use the *p*-curve model’s posterior probability as the test statistic for a permutation test, and thus obtain a frequentist *p*-value for prevalence or effect size differences, though we think this is not usually necessary.

### Example A: Interoception

In a previous study, we used a custom mixture model to estimate the population prevalence of those who can feel their own heart beating ^16^. Participants saw two circles pulsing, side-by-side, on the screen in front of them; one circle was synchronized to their cardiac systole (the phase in which blood pressure increases following a heart contraction, triggering baroceptors in the arteries) and the other pulsed exactly anti-phase. Participants were asked, on each trial, to guess which circle was synchronized to their heartbeat sensations.

In this previous study, we originally modeled the number of correct trials *k_i_* for each participant *i* as *k_i_* ∼Binomial(*n*_trials_, 0.5) under *H*_0_ and as *k_i_* ∼Binomial(*n*_trials_, *k_i_*) for some accuracy *k_i_* ∈ (0.5,1.0) under *H*_0_, and we sampled from the posterior of that mixture model given the observed *k*_H_’s from 54 participants to obtain an estimate of the prevalence of *H*_1_ in the sampled population (i.e. above-chance interoceptive accuracy). The posterior expectation for the prevalence of cardiac perceivers was 0.11 (95% HDI: [0.01, 0.21]). This model is visualized in Figure 6a.

**Figure 6:**
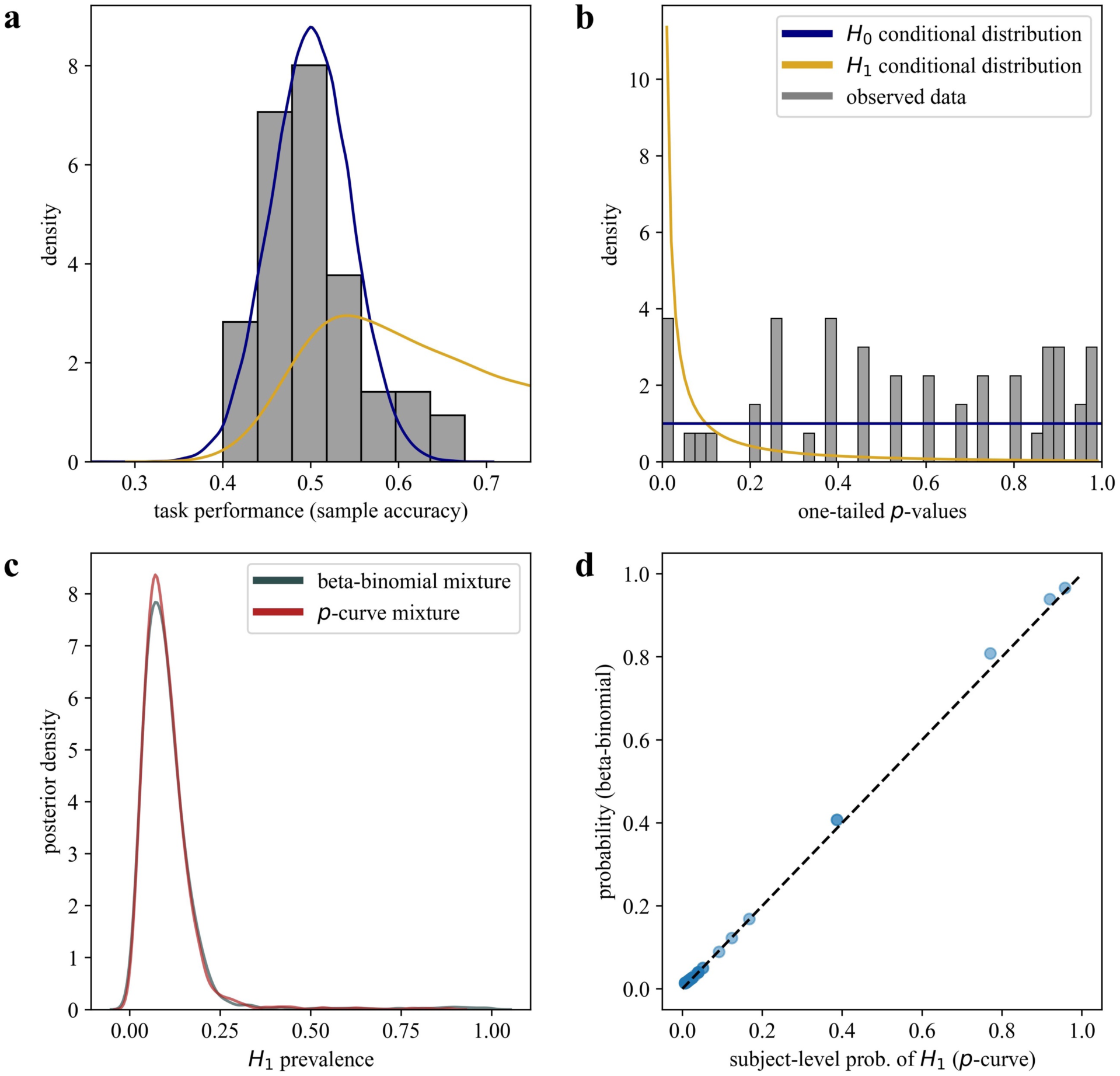
Null hypothesis significance tests as transformations of variables. (a) In a previous study (n = 54), we estimated the prevalence of above-chance performers on a discrimination task by modeling the distribution of participants’ accuracies as a mixture between a Binomial distribution for participants for whom *H*_0_ was true (at-chance accuracy) and a Beta-Binomial distribution for participants in which *H*_1_ was true (above-chance accuracy). (b) We could have instead converted the accuracies to *p*-values using a binomial test and modeled the distribution of *p*-values as a mixture between two *p*-curves. (c) Both models result in almost identical posterior distributions for the population prevalence and (d) identical per-participant posterior probabilities of *H*_1_.

Now, with *p*-curve mixture models, we can estimate the same quantity without constructing and programming a custom model. Converting the participants’ accuracies to *p*-values using a one-tailed binomial test and then plugging those *p*-values into the *p2prev* package yielded a nearly identical posterior (M = 0.10, 95% HDI: [0.01, 0.20], see Figure 6c). It is also possible to estimate the per-participant probability of *H*_1_ in both models – this was, in fact, what we were actually trying to do in the original study – and we find even these per-participant probabilities are almost the same in the two models (see Figure 6d). Still, it is worth noting one disadvantage of the *p*-curve model: the custom mixture model yields an effect size estimate in units of accuracy, while the *p*-curve model only gives an uninterpretable, unitless effect size (although that effect size parameter can be converted into a statistical power at a chosen significant level).

### Example B: Pitch Perception

Absolute pitch (AP) – the ability to recognize musical notes by their pitch alone -- is generally cited as quite rare with only 1 in 10,000 demonstrating this; even trained musicians normally require the aid of a reference note ^40^. A common claim in the literature is that experience speaking a tonal language increases the likelihood that one will develop absolute pitch. However, studies tend to dichotomize participants into AP and non-AP groups based on whether they exceed some threshold accuracy. The number of participants who exceed any binary threshold, regardless of whether it was set arbitrarily or by some statistical criteria, is in principle a function not just of the prevalence of AP but also of the pitch-labelling accuracy of those participants who do have AP. Thus, an alternative explanation for a higher proportion of tonal language speakers clearing some threshold is an increase in effect size, not in prevalence.

In a previous study ^15^, we collected behavioral judgments in a pitch-labeling task from a large sample of online participants – many of whom were attracted, unconventionally, by an article in the Wall Street Journal ^41^. Here, we analyze that dataset to estimate the difference in AP prevalence and in within-participant effect size. We first input the one-tailed *p*-values given by a binomial test on participants’ accuracies into *p2prev* to fit a *p*-curve mixture model on the full sample, estimating a prevalence of 0.53 (95% HDI: [0.46, 0.61]). It is important to note that prevalence estimates are always for the *sampled population,* which is usually not the general population. Here, our participants opted to take an online quiz to see if they have AP after reading about it in a newspaper article, so our prevalence estimates likely refer to a population that has self-selected for believing they are above-chance at naming musical notes.

Then, we fit *p*-curve mixture models to tonal language speakers and other participants separately, and we subtract posterior samples between groups to approximate the posterior of the difference (see **Methods: Between-group differences**). This results in a 95% highest density interval of [-0.02, 0.35] for the prevalence increase as a result of speaking a tonal language – not evidence against a prevalence increase by any means but the HDI does still contain zero as a plausible difference. Interestingly, however, the HDI for the within-participant effect size substantially departs from zero (95% HDI: [0.18, 1.22]), which is fairly strong evidence that tonal language experience increases the within-participant effect size *given that a participant already has AP*, seemingly in contrast to how the effect of tonal language is framed in the literature.

Of course, the dataset on which we did this analysis is quite idiosyncratic, so we do not mean to suggest that the AP literature should reevaluate its canon based merely on Example B in a methods paper. However, while our **Introduction** and simulation results warn against interpreting differences in the mean effect size as effect size differences per se, this example nicely illustrates that putative prevalence differences may also turn out to be accounted for by effect size differences when subjected to additional statistical scrutiny. Theoretical assumptions should always be evaluated explicitly, and *p*-curve mixture models provide a broadly applicable tool for doing so.

### Example C: Precision fMRI

It is becoming increasingly common in the neuroimaging literature to collect lots of data from very few participants, rather than a bit of data from many participants as in a traditional group study. This “precision neuroscience” approach allows one to detect within-participant effects that would wash out in a group average due to poor spatiotemporal alignment across participants or other idiosyncrasies in the functional organization of the brain ^42^. However, as such studies tend to forgo group-mean inference in favor of within-participant statistics, the extent to which results should be expected to generalize to the population is usually left to be inferred by the reader. While some researchers have suggested effects strong enough to be observed in a single participant should be assumed to be nearly universal ^43^, this inference is contradicted by the empirical observation that large effect sizes tend to be associated with *more* rather than less heterogeneity ^2^. Such generalization claims would be substantiated empirically if explicitly support using prevalence statistics and inference. Indeed, since these densely sampled studies already perform significance testing within each participant – just as required for a *p*-curve mixture model – some researchers have already proposed that prevalence statistics are the best way to combine results across participants ^44^.

In a recent study, we collected 100 minutes of fMRI data (as part of a longer experiment) from 4 participants while they performed a motor task in an MRI scanner and we recorded their hand movements via motion tracking ^45^. We attempted to predict participants’ continuous brain activity from the internal representations of a computational model that performs the same motor task in a biomechanical simulation, aiming to approximate the “inverse kinematic” computation required to generate muscle movement activations that will move the hand to a target position. We obtained out-of-sample *R*^2^ values from this theoretically-motivated model and from a control model, and we compared *R*^2^’s non-parametrically using threshold-free cluster enhancement to obtain a familywise error rate (FWER) corrected *p*-value for every voxel in cortex ^46^. As suggested by Ince and colleagues, the lowest FWER-corrected *p*-value across all voxels can be used as a “global” *p*-value for the presence of an effect anywhere in the brain ^14^. Similarly, the lowest FWER-corrected *p*-value in a region of interest is a *p*-value for the presence of an effect in that region (with FWER correction within that region, such that the null distribution of the smallest FWER-corrected *p*-value is uniform, not across the whole brain which would result in a right skewed distribution). This allows researchers to abstract over spatial misalignment when aggregating results across participants, since they can ask “what is the prevalence of within-participant effects across all of X area” rather than “is there a group-mean effect in any voxel in X area.”

The smallest FWER-corrected p-values across cortex for each participant were 0.00060, 0.02999, 0.04939, and 0.94601, so we could reject the null hypotheses that our theoretical model does not outperform the control model in ¾ participants at significance level α = 0.05 or in ¼ participants at significance level α = 0.01. Consequently, when we applied the Binomial prevalence model, we obtained totally different estimates depending on whether we used α = 0.05 (prevalence = 0.721, 95% HDI: [0.382, 1.00]) or α = 0.01 (prevalence = 0.504, 95% HDI: [0.114, 0.993]). Our discomfort with this estimator’s dependence on an arbitrary parameter is what motivated us to develop *p*-curve mixture models in the first place. When we put our *p*-values into *p2prev,* we estimated a prevalence of 0.610 (95% HDI: [0.224, 0.972]). As seen in Figure 3, the 0.610 posterior expectation is likely not very meaningful at such a small sample size; however, the HDI bounds maintain strong false coverage rate properties even at very low sample sizes and are thus informative.

While three of our participants had low within-participant *p*-values, one participant had a much higher *p*-value. Is this enough evidence to suggest that the participant showed no effect, or is it the case that we were just underpowered to detect it? Do we even have enough evidence to support that our participants with low *p*-values do show an effect or were they statistical flukes? *P*-curve models can help with this as well, as we can compare our mixture model to *p*-curves for the null and alternative hypothesis alone. When we calculated Bayes Factors (see **Methods: Bayesian Model Comparison**), we obtained BF = 42.03 when comparing the mixture model vs. the *H*_0_-only model, indicating that the observed data were more than 42 times more likely under the mixture model than under the model in which the null hypothesis is always true. This is strong evidence that our participants indeed show an effect. When we compare the mixture model to the *H*_1_-only model, on the other hand, we obtained a much lower Bayes Factor of BF = 3.86. While evidence leans in favor of the mixture model, indicating that a model in which not all participants express the effect can explain the high *p*-value better than the *H*_1_-only model, the evidence is not resounding. (For reference, some journals require Bayes Factors of at least 10 to support claims, though this is somewhat arbitrary). Of course, evidence should not be resounding; we only saw one low *p*-value! Nevertheless, we obtain informative results even with a small sample size, and Bayesian model comparison provided a rigorous way to collectively evaluate our within-participant results without requiring us to use an arbitrary α threshold.

### Example D: EEG Decoding

In a recent study with 25 participants, we used functional electrical stimulation (FES) of arm muscles to usurp participants’ intentional motor control in a response time task ^47^. By carefully timing the latency of FES to preempt participants’ volitional movements, we were able to elicit electrically-actuated finger movements that, under controlled timing conditions, participants either claim they themselves caused the movement or that they did not cause the movement. We accomplished this using an adaptive procedure that, for each participant, estimated and simulated at the FES latency at which participants responded – when asked after each trial whether they or the FES caused the button press in the reaction time task. The timing was set so that that participants reported that they caused the movement on roughly 50% of trials. This approach created balanced sets of trials participants perceived as self- or other-caused, so that their sense of agency (SoA) could be decoded from their EEG.

While we reported group-averaged results in the original study ^47^, as is standard, in analysis of the data we noted substantial heterogeneity in how participants responded to the adaptive procedure. In some participants, the adaptive procedure honed in on their threshold early and FES remained at that threshold latency for the rest of the experiment. In these participants, a within-participant logistic regression predicting agency judgments from FES latency would return significant results, as participants were highly sensitive to deviations from their threshold. However, other participants’ thresholds seemed to slowly shift over time throughout the experiment; in these participants, the same logistic regression would yield non-significant results, as the same 50/50 response distribution was observed across a range of latencies. With *p*-curve mixture models, we can now better assess how decoding of agency judgments from EEG is related to this behavioral difference.

Using a within-group prevalence model, we can model the joint probability of participants’ agency judgments being sensitive to FES latency around their threshold (i.e. are “stable-threshold participants” vs. “unstable-threshold participants”) and of agency judgments being decodable from their EEG – for which we computed a *p*-value for the 10-fold cross-validated decoding AUROC using a permutation test at each time-point in the epoch following FES onset. As described by Ince et al. (2021), we can perform prevalence inference at each time in the EEG epoch, or we can take the lowest familywise error rate corrected *p*-value across participants as a “global” *p*-value testing the null hypothesis that agency judgments are not decodable from their EEG at any point. Familywise error rates corrected p-values were compute using the maximum statistic method ^48^. Using the lowest familywise error rate corrected *p*-value in this way satisfies the assumptions of the *p*-curve mixture model, as its null distribution is uniform, but this would not necessarily be the case for, say, a false discovery rate corrected *p*-value.

Using the within-group *p*-curve model described in **Methods**, we estimate the prevalence of behavioral sensitivity of agency judgments to FES latency – over the whole epoch – as 0.655 (95% HDI: [0.473, 0.870]) and that of decodable agency judgments as 0.922 (95% HDI: [0.831, 0.996]), with evidence for a prevalence difference of 0.268 (95% HDI: [0.054, 0.479]). Logically, as the prevalence of decodable agency judgments exceeds that of the behavioral sensitivity effect, we can already conclude that above-chance decoding performance can be achieved for both behavioral phenotypes. However, our within-group prevalence model also allows these conditional probabilities to be estimated explicitly. The prevalence of EEG-decodable agency judgments among stable-threshold participants is estimated as 0.942 (95% HDI: [0.835, 1.000]) and among unstable-threshold participants is 0.881 (95% HDI: [0.668, 1.000]), without strong evidence for a difference (0.061, 95% HDI [-0.169, 0.321]). However, just because there is not a difference in prevalence *across the whole EEG epoch* does not mean there is no difference at all. We can also calculate the conditional prevalences across time as seen in Figure 7.

**Figure 7:**
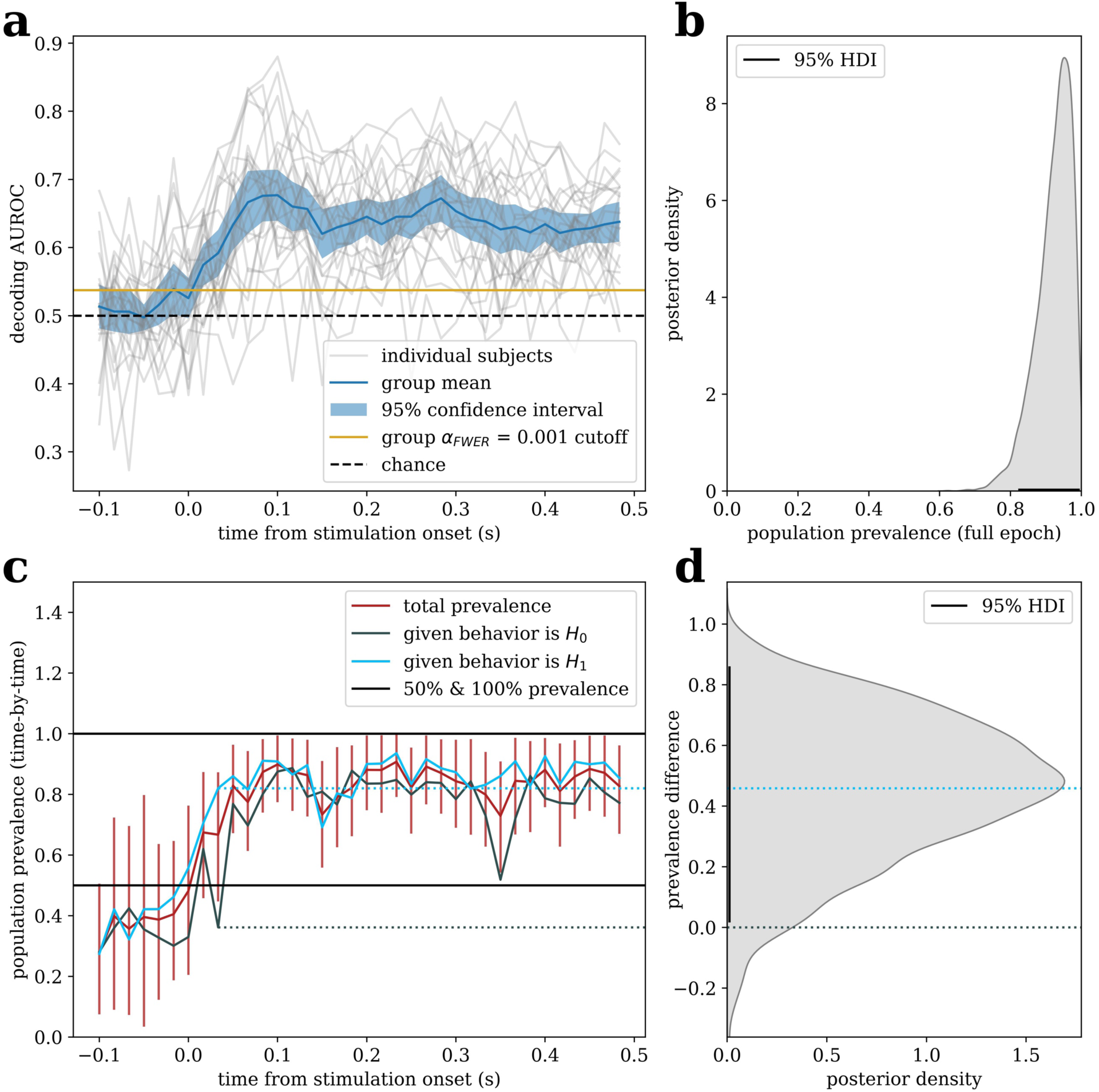
Prevalence estimation across time with EEG decoding results. (a) The group-mean and single participant decoding performance time courses for predict agency judgments from the EEG as described in Example D (n = 23). (b) The posterior for the population prevalence of decodable agency judgments at any point in the epoch. (c) The prevalences of decodable agency judgments at specific times in the EEG relative to electrical muscle stimulation, with posterior means for both total prevalences and prevalences conditional on the behavioral effect, and 95% HDIs for total prevalence. Note that posterior HDIs for prevalence never include zero, as prevalence is *a priori* in the interval [0, 1]; thus, prevalence estimates should only be interpreted when there is other evidence of an effect from Bayesian model comparison or from an appropriate group-level test. (d) The posterior prevalence difference conditional on the behavioral effect at 0.033 seconds, showing stable-threshold participants have a higher prevalence of decodable information at this time.

As seen in Figure 7c, the prevalences conditional on being a stable-threshold or unstable-threshold participant can be computed at each time following stimulation, and a posterior difference can be estimated. For example, at 0.033 seconds after stimulation, just after the initial cortical response – which is expected at around 20 milliseconds for stimulation on the wrist ^49^ – decodable information is already present in a majority of stable-threshold participants (0.820, 95% HDI: [0.588, 1.000]). However, unstable-threshold participants are less likely to show decodable information at this time point (prevalence = 0.361, 95% HDI: [0.022, 0.688]; difference = 0.458, 95% HDI: [0.016, 0.861]). In other words, the neural responses to muscle stimulation tend to predict agency judgments earlier in stable-threshold participants, speculatively reflecting increased sensitivity to low-level sensorimotor mismatches.

Notably, while it is possible that bifurcating participants based on their judgment stability might allow group-mean approaches to detect a behavior-contingent difference in *mean* decoding performance, the prevalence estimation approach is able to support the claim that decodable information is *absent* at this time in a higher proportion of unstable-threshold participants. As information-based measurement such as decoding accuracies are known to show significantly above-chance group-mean performance even when decoding is only achievable in a minority of participants ^12^, it is unlikely group-mean approaches – even those that can support null results in principle, such as equivalence tests or Bayes factors – could ever support such a finding unless prevalence were actually zero. Indeed, for this same reason, significant group-mean decoding results actually do not generally support the claim that decoding is possible in a majority of participants; formal prevalence inference is required to assess the population generalization of any “multivariate pattern analysis” (MVPA) study, though this fact is frequently ignored in the literature ^12,50^. This can be seen in Figure 7, however, as significant group-mean decoding performance precedes the time at which we can conclude that prevalence exceeds 50% of the population. Conversely, the group-averaged time courses in Figure 7a may give the impression that decoding time courses are smooth, but there are numerous instances where we can conclude the prevalence of decodable information is well below 100% in Figure 7c. This indicated that individual decoding time courses are heterogenous, which is not at all obvious from looking at the single participant time courses and could easily be dismissed as noise around a population mean.

Note that the HDIs for prevalence estimates in Figure 7c exclude zero even when mean prevalence is at chance. This is guaranteed to occur in practice, as prevalence is bounded to the interval [0, 1] and thus no posterior probability mass can fall below zero. Thus, it is important to remember that prevalence estimates are only meaningful when there is sufficient evidence against the all-*H*_0_hypothesis being true; such evidence can be quantified as described in **Methods: Bayesian Model Comparison.**

Finally, as out-of-sample prediction in common cross-validation schemes are not identically and independently distributed due to dependence across folds ^51^, and thus decoding performance metrics tend not to follow a known distribution, performing power analyses for MVPA studies is challenging even among other neuroimaging studies. As *p*-curve effect size estimates, though uninterpretable in-and-of-themselves, can be easily converted into within-participant power estimates at any chosen significance level (see **Methods: The *p*-curve mixture model**), we can report within-participant power estimates *post hoc,* which can be extremely useful for planning future research. In this example, the within-participant power for detecting above-chance decoding at any point over the EEG epoch at significance level 0.05 is 0.956 (95% HDI: [0.918, 0.988]). Note that, in addition to providing a valid point estimate, our approach yields a credible interval (or full posterior) for the within-participant power; in contrast, *post hoc* power estimates for group-mean NHSTs are essentially useless, providing no information beyond that given by the *p*-value itself ^52^.

## Discussion

### Epistemology of statistical inference

We imagine that the use of *p*-values within a Bayesian analysis may surprise some readers, though this practice is not unprecedented ^14,28^. It is important to note that the use of *p*-values does not entail adoption of the frequentist epistemology within which *p*-values are most often seen. *p*-values are, simply put, a transformation of data from original units to units in which independent observations have the Uniform(0,1) likelihood under some well-defined (albeit sometimes fictional) “null” distribution; this transformation of variables is derived from probability theory, which deals with probabilities in the abstract, agnostic to any particular statistical epistemology (e.g. Bayesian or frequentist) that postulates how probabilities should be interpreted.

In the context of the frequentist epistemology, probabilities are simply long-term rates (e.g. the rate at which a fixed parameter will fall within an interval if that interval is recomputed across many repeated random samples). In such a framework, it is logical to perform inference by subjecting *p*-values to a binary threshold at a desired false positive rate. But that thresholding is how *p*-values are *used* in the context of null hypothesis significance testing (NHST) rather than what they *are.* Analogously, Bayes’ theorem – the rule for inverting conditional probabilities – is derived from probability theory and often used within frequentist statistics (for instance, to calculate the probability/*rate* at which the alternative hypothesis will be true when significant results are obtained from a medical test given the test’s error *rates* and the base *rate* of a disease). Within a Bayesian epistemology, however, probabilities are not merely rates but represent *beliefs* (and their uncertainty) which may be updated given fixed observations of data. In Bayesian statistics, then, Bayes’ theorem takes on additional meaning as the optimal rule for belief updating. Our Bayesian approach uses *p*-values but, rather than thresholding them as in NHST, treats them as fixed observations used to update beliefs about parameters (prevalence and effect size).

### Incorporating Prevalence Estimation into the Literature

Behavioral scientists tend to report experimental findings as if they should be expected to pertain to all of, or at least the majority, of the population from which they recruited their participants. Usually, these claims are supported by a statistically significant group-level effect size measurement. However, as noted in the Introduction (see Fig. 1) nonzero effects measured at the group level do not imply that a majority of participants in the sample, much less in the population, show that effect. Indeed, claims about the typicality of an effect require explicit statistical support, such as by directly estimating the population prevalence.

When we began developing *p*-curve mixture models, we were anticipating they would be applied primarily to dense sampling studies, offering only modest benefits over prevalence estimation methods that require the power of within-participant tests to be near 100% ^14^. It is important to note, however, that we obtain strong performance even when (simulated) participants have fairly low trial counts; 50 trials per condition in an EEG as experiment, as in our simulations, is certainly quite modest. Moreover, we were surprised to learn that *p*-curve mixture models can, in some cases, be applied without any loss in sensitivity compared to group-level null hypothesis significance testing, as in our within-group difference simulations where we even saw sensitivity *gains* (see Figure 5c). Applying these models instead of, or in addition to, traditional NHST can provide a more complete description of effect distributions within and across populations – a critical tool, we believe, for a time in which population heterogeneity in both psychological traits and functional brain organization are increasingly discussed ^10,42^.

Indeed, a strong limitation of existing methods for prevalence estimation, save for Bayesian mixture modeling ^16^, has been that they only provide lower bounds on the population prevalence ^12,14^, or otherwise require nearly 100% statistical power to yield accurate estimates of the true prevalence ^14^. As such, recent efforts toward prevalence estimation in the behavioral sciences has been primarily geared toward studies explicitly designed to achieve very high within-participant power ^44^. Bayesian mixture models can provide an estimate of prevalence regardless of the within-participant power of a study and can thus be applied to datasets from experiments originally designed to test for differences in group means. This wide applicability allows researchers to easily quantify, with appropriate uncertainty, the proportion of the population to which their findings are expected to represent. This metric may provide crucial insight into how well-suited basic science findings are for translation to clinical settings, and empirical prevelance measurements could shed light on the causes of non-replications as researchers debate the role of generalizablity in precipitating the replication crisis ^2,4–7^.

## Limitations

Nonetheless, *p*-curve mixture models have some substantial limitations that potential users should take under advisement.

Firstly, our approach assumes the existence of truly null effects, which may not always be appropriate or meaningful. Many researchers are used to such an assumption, as it is the same assumption adopted in null hypothesis significance testing procedures (i.e. the “null hypothesis” itself). The assumption is not uniquely frequentist; Bayes Factor approaches to statistical inference often consider a null model with an effect size of exactly zero as one of several competing explanations for the observed data ^20,53^. However, some statisticans argue that an effect size of exactly zero (or whatever other null value) is not a reasonable possibility for continuous parameters ^54^; in this view, the null hypothesis is no more than an occassionally “useful fiction.” As one of our reviewers pointed out, it may not be appropriate to elevate such a fiction to the level of an ontological claim that there is indeed a qualitiatively distinct subgroup of participants who do not show an effect at all unless there is very good reason to do so. Under such a view, estimating the prevalence of an would make little sense. We encourage readers to read the open reviews of this paper for a more in depth discussion of this issue, as well as Rouder and Haaf’s recent review of theoretical issues pertaining to the possibility of qualitative (rather than quantitative) individual differences in cognition ^55^. The present authors believe that a null hypothesis is, while perhaps a fiction, indeed useful in many cases. However, we agree that the assumption that there are subgroups of subjects within a population (e.g. those that show an effect and those that do not) should not be made without evidence. Prevalence estimates should always be accompanied by support for the implied heterogenity structure (i.e. a mixture of distributions as opposed to a single distribution), which we suggest can be accomplished using Bayesian model comparison as described in **Methods**, and users should always exercise their own judgment with respect to the *a priori* assumptions underlying their analysis approach.

Researchers should also note that *p*-curve mixture models make the assumption that the distribution of repeated, independent *p*-values is uniform under the null hypothesis. This is, of course, to say that users should select an appropriate procedure to compute *p*-values. Anytime the assumptions of that procedure are violated in a way that would result in over- or under-conservative inference in a null hypothesis significance test, *p*-curve mixture models are also likely to yield poor estimates. Even non-parametric approaches to computing *p*-values carry assumptions of which researchers must be mindful. Moreover, tests for some types of data only produce a uniform distribution of *p*-values under the null distribution asymptotically; for example, a binomial test on discrete data can only produce a finite number of *p*-values, so the distribution of *p*-values is only approximately uniform for finite trial counts. Notably, we did use discrete data (sample accuracies) with a relatively modest trial count in the simulations shown in Figures 3-4, and our approach seemed robust to that particular violation of assumptions. However, researchers should always exercise caution during analysis, especially in cases where observations are not continuously valued and trial counts are very small.

It should also be kept in mind that prevalence estimates only pertain to the population from which the study sample was drawn; if a sample is (systematically) unrepresentative of the population to which a researcher intends to generalize, then the prevalence estimate will not be valid. (For example, results from Example B, where sampling was not random, should not be expected to extend to a general population) In that case, prevalence inference is best limited to formal comparison between all-*H*_0_, all-*H*_1_, and mixture models of the data; it may still be possible to rule out the all-*H*_0_ (nobody shows effect) and all-*H*_1_ models (everyone shows the effect) even from an unrepresentative subsample. This could conceivably be accomplished with *p*-curve mixture models (see **Methods: Bayesian Model Comparison**), but we also encourage researchers to consider other approaches that may depend on less restrictive assumptions when full prevalence inference is not being used ^3,20^.

In addition, *p*-curve mixture models treat all participants who show an effect as coming from the same distribution. Thus, they would not detect a case in which some participants show an effect in the opposite direction of the group average; while this scenario may sound unlikely on the surface, this has been shown to occur in commonly used experimental tasks ^3^. Moreover, metaanlaytic findings seem to suggest that some fields of behavioral science – such as brain training, video gaming, mindset, and stereotype threat research – are particularly prone to produce treatment effect distributions with multiple modes ^56^. While our approach seems to be robust against overdispersion under the alternative hypothesis (see Fig. 4), and thus we would not expect prevalence estimates to be wrong in this case per se, a *p-*curve mixture model would fail to characterize this important feature of the distribution of participant-level effect sizes. In this vein, while quantifying prevalence is an easy practice for empirical researchers to adopt in the interest of improving the precision and scope of their claims, it may only be the first step required to assess generalization in a research program aiming to design meaningful interventions or influence public policy ^9^.

Lastly, in a case where many such hypotheses are tested in parallel such as in Example D or most modern neuroimaging settings, we suggest at least reporting conventional familywise-error rate corrected NHST results (as in Fig. 7a) alongside prevalence estimates if results are being used to draw binary conclusions about the existence of an effect – or, even better, a complementary approach to prevalence inference that explicitly accounts for multiple comparisons across sensors, voxels, or time ^12^. Though Bayesian inference does not suffer from familywise error rate inflation due to multiple comparisons in principle (as a “fully” Bayesian approach would never draw a binary conclusion thus not produce any errors to being with), it may still lead to erroneous conclusions in practice when a huge amount of Bayesian models are fit in parallel (i.e. one model per timepoint or voxel) and used to draw a binary conclusion (i.e. there is or is not an effect). Another, fully Bayesian approach would be to aggregate all data into a single hierarchical model that appropriately pools information across times, voxels, et cetera to calibrate estimates, but such models are well beyond the scope of the present resource ^57,58^.

### When to use *p*-curve Mixtures or Other Approaches

As we illustrate in Example A (see Fig. 6), *p*-curve mixture models may in many cases yield essentially identical results to more traditional Bayesian mixture model specifications ^16^. (Indeed, the *p*-curve mixture model *is* a traditional Bayesian mixture model, specified using the likelihood of the observed data’s *p*-values rather than of the data on its original scale. This is essentially just a transformation of variables.) When a more straightforward Bayesian mixture model can be implemented, however, we would still recommend the more traditional approach, which retains several advantages over the *p*-curve mixture – notably, exact likelihoods can potentially be used, prior information may be easier to incorporate, and the within-participant effect size can be estimated in interpretable units. Alternatively, if only the question of whether some or all participants in a population show some effect, and the exact prevalence is not of interest (or unidentifiable e.g. if participants are not randomly sampled from the population the experimenter wishes to generalize to), then comparison of Bayesian hierarchical models with sign constraints can be used to evaluate whether participants all show an effect size with the same sign ^20^, often with weaker assumptions than the approach shown here. However, it may be many decades before the modal researcher has received training to craft a bespoke Bayesian model, let alone to implement it in code (which often involves arcane reparameterization tricks to enable posterior sampling) and to evaluate its efficacy in simulation. In the meantime, our model and its validated software implementation can often achieve comparable results to bespoke mixture models with little-to-no additional code; *p*-curve mixtures may not always be the best prevalence estimator for any particular case, but should perform adequately across a wide range of use cases.

In some cases, including many neuroimaging applications, it may be impractical for even seasoned Bayesian modelers to craft a bespoke model. It is no small feat to write a valid likelihood for cluster-based statistics as in our EEG simulations or in Example C even when possible in principle ^59^, or for cross-validated performance measures as in Example D. Moreover, the consensus in the neuroimaging literature is that typical parametric assumptions lead to misleading inferences in such high-dimensional or otherwise non-standard inference settings ^60,61^. Since the *p*-values used as observed data for our model can be computed non-parametrically, our approach allows researchers to proceed with weak parametric assumptions in otherwise problematic settings, as was empirically effective in our simulations (e.g. see Fig. 5). On this point, it is worth emphasizing that our estimation procedure will only be effective when *p*-values are correct (i.e. the same assumptions that would be required for a NHST are satisfied), so we encourage researchers to compute *p*-values using a non-parametric approach (e.g. permutation test) when they cannot justify the assumptions of a parametric approach. We do, however, note that even non-parametric procedures may sometimes not produce an approximately uniform distribution of *p*-values under the null hypothesis (e.g. a permutation test when the test statistic is discrete). We encourage users to check this assumption (e.g. inspect the permutation null distribution of that test statistic) whenever in doubt.

When researchers wish merely to conclude that the population prevalence exceeds a certain threshold (e.g. an effect is present in the majority of subjects), and do not care about estimating the specific prevalence per se, they may wish to use a fully non-parametric approach such as the frequentist tests previously proposed in the neuroimaging literature ^12,50^; these procedures allow for (albeit more limited, yielding only a lower bound) prevalence inference with weaker assumptions than are made by *p*-curve mixture models and, as a bonus, are extremely well suited to a multiple testing setting. Similarly, if only a lower bound on the prevalence is needed, or if the researcher is confident within-subject power is nearly 1.0 or otherwise knows the within-subject power of their design with high confidence, then the implementation of their binomial method provided by Ince and colleagues should perform well with the lowest computational cost out of all the approaches listed ^14^, again with somewhat weaker assumptions than our own approach.

## Conclusion

Many subfields in the behavioral and biological sciences carry unchecked assumptions about what their group mean differences really, well, *mean*. By providing a user-friendly interface to *p*-curve mixture models with our *p2prev* software package, we hope to facilitate the use of Bayesian mixture models so that scientists can rigorously evaluate these assumptions. Doing so need not necessarily entail collecting new data or adjusting experimental designs; our method is well-suited to existing datasets – as illustrated by some of the worked examples in this paper. As such, we hope *p*-curve mixture models work their way into the experimentalist’s toolkit.

## Data Availability

The raw datasets used in our examples are available at the open data repositories documented in the papers in which these data were originally reported ^15,16,45,47^. However, we also provide smaller, preprocessed versions of these datasets sufficient to replicate our examples in the same Zenodo repository ^62^ as our code (https://zenodo.org/doi/10.5281/zenodo.11459064).

## Code Availability

Our GitHub repository (https://github.com/john-veillette/p2prev) contains the source code for the *p2prev* package, tutorial examples including the examples in this paper, code to reproduce our simulations, and a link to the most up-to-date documentation for the package interface. The GitHub repository will be continually updated while we make improvements to *p2prev* and its documentation, but all stable releases of the *p2prev* source code and accompanying tutorial code are permanently archived on *Zenodo,* with a digital object identifier (DOI) that always leads to the most recent release (https://zenodo.org/doi/10.5281/zenodo.11459064) as well as a unique DOI for each release (e.g. the version used for the simulations in this report, *p2prev* v0.0.2). The *p2prev* package can also be installed from the PyPI package server using Python’s “pip” command. Additionally, a graphical interface for basic functionality is available online (https://jveillette.shinyapps.io/p2prev/).

## Author Contributions

**H.C.N.:** Funding acquisition, Resources, Supervision, and Writing - review & editing. **J.P.V.:** Conceptualization, Formal analysis, Funding acquisition, Investigation, Methodology, Project administration, Software, Validation, Visualization, Writing - original draft, and Writing - review & editing.

## Acknowledgements

This project was supported by NSF BCS 2024923 to H.C.N.; J.P.V. was supported by NSF DGE 1746045. This work was completed in part with resources provided by the University of Chicago’s Research Computing Center. The funders had no role in study design, data collection and analysis, decision to publish or preparation of the manuscript.

